# An alternative mechanism for skeletal muscle dysfunction in long-term post-viral lung disease

**DOI:** 10.1101/2022.10.07.511313

**Authors:** Ryan A. Martin, Shamus P. Keeler, Kangyun Wu, William J. Shearon, Devin Patel, My Hoang, Christy M. Hoffmann, Michael E. Hughes, Michael J. Holtzman

## Abstract

Chronic lung disease is often accompanied by disabling extrapulmonary symptoms, notably skeletal muscle dysfunction and atrophy. Moreover, the severity of respiratory symptoms correlates with decreased muscle mass and in turn lowered physical activity and survival rates. Previous models of muscle atrophy in chronic lung disease often modeled COPD and relied on cigarette smoke exposure and LPS-stimulation, but these conditions independently affect skeletal muscle even without accompanying lung disease. Moreover, there is an emerging and pressing need to understand the extrapulmonary manifestations of long-term post-viral lung disease (PVLD) as found in Covid-19. Here, we examine the development of skeletal muscle dysfunction in the setting of chronic pulmonary disease using a mouse model of PVLD caused by infection due to the natural pathogen Sendai virus. We identify a significant decrease in myofiber size when PVLD is maximal at 49 d after infection. We find no change in the relative types of myofibers, but the greatest decrease in fiber size is localized to fast-twitch type IIB myofibers based on myosin heavy chain immunostaining. Remarkably, all biomarkers of myocyte protein synthesis and degradation (total RNA, ribosomal abundance, and ubiquitin-proteasome expression) were stable throughout the acute infectious illness and chronic post-viral disease process. Together, the results demonstrate a distinct pattern of skeletal muscle dysfunction in a mouse model of long-term PVLD. The findings thereby provide new insight into prolonged limitations in exercise capacity in patients with chronic lung disease after viral infections and perhaps other types of lung injury.

## INTRODUCTION

Respiratory disease particularly in the form of chronic obstructive pulmonary disease (COPD) has emerged as a leading cause of morbidity and mortality (8, 31). In addition, infection with the respiratory virus SARS-CoV-2 has added a top cause of death known as coronavirus disease 2019 (Covid-19) (22) and an associated chronic disease known as long Covid (2, 9, 18). In each of these conditions, a dominant pathology stems from lung injury, inflammation, and remodeling, but there is also a systemic process with significant extrapulmonary manifestations. For example, COPD patients characteristically exhibit weight loss accompanied by skeletal muscle dysfunction with marked impairment of muscle strength and endurance (10, 13, 16, 21, 29). This dysfunction is manifested primarily as loss of muscle mass, also known as muscle atrophy, as well as a phenotypic shift in myofiber composition. Specifically, peripheral muscle transitions from a mixture of oxidative and glycolytic myofibers to a predominant glycolytic phenotype (14, 19), signifying a change from Type I and Type IIA (slow and fast oxidative fibers) to Type IIB (fast glycolytic) fibers that primarily use anaerobic glycolysis (14). Patients with COPD-related muscle dysfunction have reduced quality of life, activities of daily living, and poorer disease outcomes (23). However, the underlying mechanisms of COPD-associated muscle atrophy remain poorly defined. Similarly, there is limited understanding of muscle phenotype in post-viral lung disease (PVLD) in general and patients with Covid-19 in particular wherein fatigue is as common as respiratory symptoms and exercise limitation is linked to skeletal muscle dysfunction (1, 2).

To better define the long-term process for skeletal muscle dysfunction in lung disease requires a relevant experimental model. Existing models include inhaled exposures to air pollutants, fumes, or tobacco smoke, which induce pulmonary disease but may confound interpretations of muscle abnormalities (32). For example, even acute cigarette smoke exposure in a mouse model causes muscle atrophy (17), reduces muscle strength (20), and alters muscle innervation in the absence of pulmonary disease (11). Moreover, the muscle dysfunction in this model reverses upon smoke cessation, whereas muscle dysfunction continues and even progresses long-term in humans with COPD (7, 33) and Covid-19 (2, 9, 18, 37). Moreover, smoke-inhalation models often cause only mild disease (32) even with long-term exposure (∼24-36 weeks) (15, 38). These key differences from the comparable human conditions underscore the need for a new animal model that better captures the features of severe and persistent lung disease using a natural trigger such as respiratory viral infection. This type of model might also prove useful for the study of skeletal muscle dysfunction in general given the broad role of viral infection for initiation, exacerbation, and progression of respiratory diseases.

Here we use this rational to engage a mouse model of chronic lung disease that develops after infection with the natural pathogen Sendai virus (SeV) (27). In this model, the animal experiences a severe acute infectious illness and then switches to life-long lung remodeling disease that includes basal-epithelial stem cell reprogramming, mucinous differentiation, immune cell activation, and airway hyperreactivity (5, 12, 27, 28, 34-36). These features as well as underlying cellular and molecular abnormalities are also found in humans with chronic lung disease due to COPD and asthma (3, 5, 6, 12) and overlap significantly with the phenotype in severe and long-term Covid-19 (37). However, the possibility that this type of respiratory viral infection might also induce long-term skeletal muscle dysfunction remained unstudied to our knowledge. Here we address this issue and find marked changes in skeletal muscle morphology with implications for patients with muscle dysfunction typical of human patients with chronic respiratory disease especially in forms triggered and/or exacerbated by respiratory viral infection.

## MATERIALS AND METHODS

### Mouse procedures

All experimental procedures involving animals were approved by the Animal Studies Committees of Washington University School of Medicine in accordance with guidelines from the National Institutes of Health. All mice were maintained and co-housed in a barrier facility using cages fitted with micro-isolator lids. Sendai virus (SeV, Sendai/52 Fushimi strain, ATCC VR-105) was obtained from ATCC, and stocks were grown in embryonating chicken eggs and titered by plaque-forming assay as described previously (12). Male C57BL/6J mice (#000664) were obtained from Jackson Laboratory and infected with SeV (2.6 × 10^5^ PFU) as described previously (35). Briefly, virus was delivered intranasally in 30 μl of phosphate-buffered saline (PBS) or an equivalent amount of PBS containing UV-inactivated virus (SeV-UV) as a control. Delivery was performed under ketamine/xylazine anesthesia using mice at 6 weeks of age. Total body weights were recorded daily to evaluate the severity of infection. At 8, 12, 21, and 49 d after SeV infection or SeV-UV control, skeletal muscle tissue was collected, placed in O.C.T. Compound (Fisher Scientific) in isopentane cooled with dry ice, and then stored at -80 °C. Additional muscle samples were collected for RNA extraction and analysis.

### Immunofluorescence microscopy

Transverse sections were air dried for 30 min and blocked with 10% bovine serum albumin (Fisher Scientific; BP9700) in PBS. Sections were washed and labeled with a combination of unconjugated rabbit anti-laminin antibody (Abcam Cat# ab11575, RRID:AB_298179) (1:400) and mouse antibody against a specific type of myosin heavy chain (MHC): Type I (DSHB Cat# BA-F8, RRID:AB_10572253), Type IIA (DSHB Cat#SC-71, RRID:AB_2147165), Type IIX (DSHB Cat# 6H1, RRID:AB_1157897), or Type IIB (DSHB Cat# BF-F3, RRID:AB_2266724) (1:10). Primary antibodies were visualized after incubation with species and isotype specific antibodies conjugated with AlexaFluor® 568, 488 IgG, or 594 IgM (1:200 dilution). Sections of each muscle labeled with one type of MHC were imaged at 200x magnification to determine myofiber type distribution. We captured up to 16 images in identical areas of each MHC^+^ section that resulted in 488 ± 184 myofibers per muscle. MHC^+^ myofibers were identified using Fiji software (ImageJ) at standard color threshold, and cross-sectional area was determined for MHC^+^ myofibers using the semi-automated macro used for histology analysis.

### Western blotting

Quadricep muscles of SeV-infected and SeV-UV-control mice were used for gel electrophoresis and Western blotting. For these experiments, muscle samples were bead homogenized by a Qiagen Tissuelyser in reducing sample buffer (Tris-HCl, pH 6.8; 2% SDS; 10% glycerol; 1% β-mercaptoethanol) containing protease inhibitors (Cell Biolabs, AKR-190). Samples were centrifuged at 10,000 rpm at 4 °C, and resulting supernatants were stored at -80 °C. Sample concentrations were determined at 280 nm absorbance using a NanoDrop Spectrophotometer. Protein samples were boiled and separated on Nu-Page 4-12% Bis-Tris gels (NP0336; Invitrogen) for 2 h at 120V and transferred to nitrocellulose membranes using the iBlot 2 mini-stacks (IB23002; Invitrogen) according to the manufacture’s guidelines. Membranes were washed with Tris-buffered saline containing 0.1% Tween 20 (TBS-T), blocked in 5% non-fat dry milk, and incubated overnight at 4° C with antibodies for MuRF-1 (Proteintech Cat# 55456-1-AP, RRID:AB_11232209) (1:1000), atrogin-1/MaFbx (ECM biosciences Cat# AP2041, RRID:AB_2246979) (1:1000) and GAPDH (Proteintech Cat# 60004-1-Ig, RRID:AB_2107436) (1:20,000). Bound antibodies were detected using species and isotype specific IRdye 680 or 800 (Licor; 1:15,000) antibodies and visualized using an Odyssey infrared scanner.

### Ribosomal RNA (rRNA) analysis

Total RNA and ribosomal content were determined in quadriceps muscle after digestion in Trizol Reagent™ using the manufacturer’s protocol. RNA content was then isolated using Qiagen RNeasy columns. After the final centrifugation, RNA samples were eluted in RNase-free water for quantification. Samples were analyzed on the Agilent BioAnalyzer using the Agilent RNA 6000 Nano Chip to determine RNA concentration and abundance of 18S and 28S rRNA. Ribosomal RNA abundance was expressed as the total 18S and 28s rRNA per mg of muscle mass.

### RNA-sequencing (RNAseq)

RNAseq was performed using an Illumina HiSeq 3000. The resulting 50-bp, single-end reads were aligned to the genome and transcriptome of *Mus musculus* (ensembl GRCm38/mm10) using STAR v 2.6.0a. Gene level TPM values were calculated using RSEM v.1.3.3.

### Statistical analysis and reproducibility

Data are presented as mean and SEM unless otherwise stated. Comparisons between groups were performed in SPSS statistical software (IBM) using a two-way Analysis of Variance (ANOVA) with time-course and treatment as factors and statistical significance set at *P*< 0.05. A Tukey post-hoc test was performed to determine any specific differences corrected for multiple comparisons. Pearson correlational analysis was performed on select outcomes with statistical significance set at *P* < 0.05. The number of mice for each condition is defined in the Figure Legends.

## RESULTS

### PVLD is linked to skeletal muscle atrophy

To first determine if our animal model of PVLD is accompanied by a change in skeletal muscle phenotype, we developed a method to identify myofibers in quadriceps muscle obtained from mice with and without SeV infection. Based on immunostaining for laminin in transverse sections of muscle tissue, we could identify muscle cell basal lamina under these experimental conditions and thereby assess muscle fiber size. Qualitative examination of immunostaining showed no change in cross-sectional area of individual myofibers at 12 d after SeV infection compared to SeV-UV controls (Fig. 1. left column). Moreover, we found a striking increase in this area from 12 to 49 d in the SeV-UV control mice (Fig. 1, top row) consistent with ongoing growth with age. However, we also observed a marked decrease in this area in SeV-infected compared to SeV-UV control mice at 49 d (Fig. 1, right column). These time points fit with the progression of PVLD that is not well developed at 12 d after SeV infection but is fully manifest at 49 d after infection based on our earlier work (35, 39). Further, the differences in myofiber size developed despite similar body weights in infected and control mice (data not shown), indicating that decreases in myofiber size were not attributable to lower body mass.

**Figure 1.**
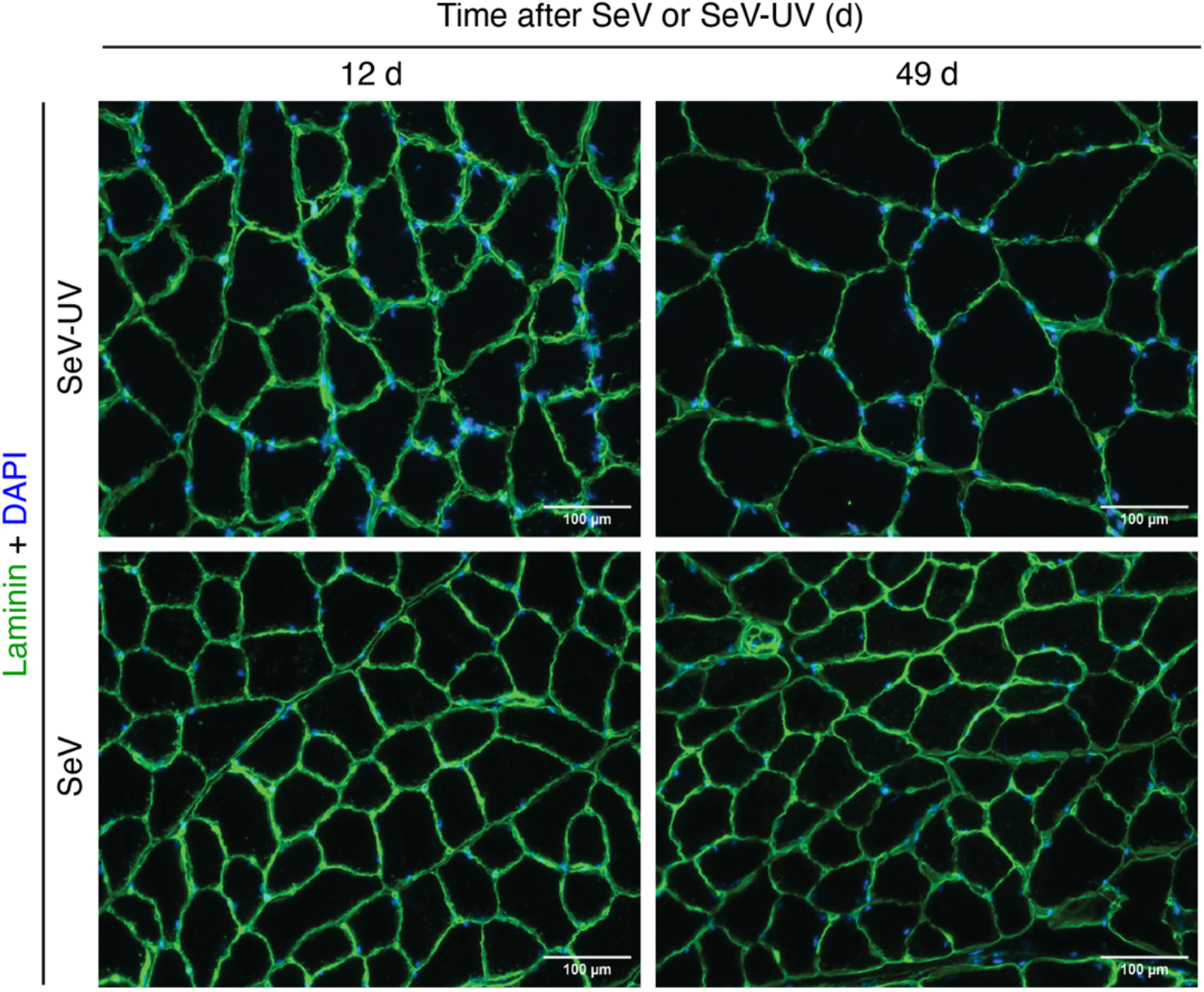
Identifying skeletal muscle myofibers in a mouse model of post-viral lung disease (PVLD). Immunostaining for laminin and DAPI counterstaining was performed in cross-sections of quadriceps muscle obtained from mice at 12 and 49 d after SeV infection or SeV-UV control condition. Basal-lamina^+^ staining was used to identify individual myofibers with associated DAPI^+^ staining nuclei. Scale bars = 100 μm. Results are representative of n = 3-6 mice per experimental condition.

To determine the significance of the observed effects on myofiber size, we rigorously quantified cross-sectional area (μm^2^) of myofibers in transverse sections (10-μm in thickness) of quadriceps muscle tissue. For power of the analysis, we assessed 1,554 ± 239 (mean ± SD) myofibers after SeV infection and 1,451 ± 304 (mean ± SD) myofibers per timepoint after SeV-UV control. Analysis of myofiber size based on cross-sectional area showed a significant increase in size at 49 d compared to 12 d after SeV-UV consistent with expected animal growth at the age of study (Fig. 2A). Moreover, myofiber size was similar in mice studied at 12 d after SeV infection or SeV-UV control (Fig. 2A). However, we detected a significant decrease in myofiber size at 49 d in SeV-infection compared to SeV-UV control mice (Fig. 2A). Similarly, our analysis of frequency distribution based on myofiber cross-sectional area showed a similar distribution of myofibers in mice at 12 d after SeV-infection or SeV-UV control whereas we found a decreased abundance of larger myofibers at 49 d after SeV-infection compared to SeV-UV mice (Fig. 2B). Our results demonstrate that myofiber growth is inhibited during the development of PVLD after SeV infection.

**Figure 2.**
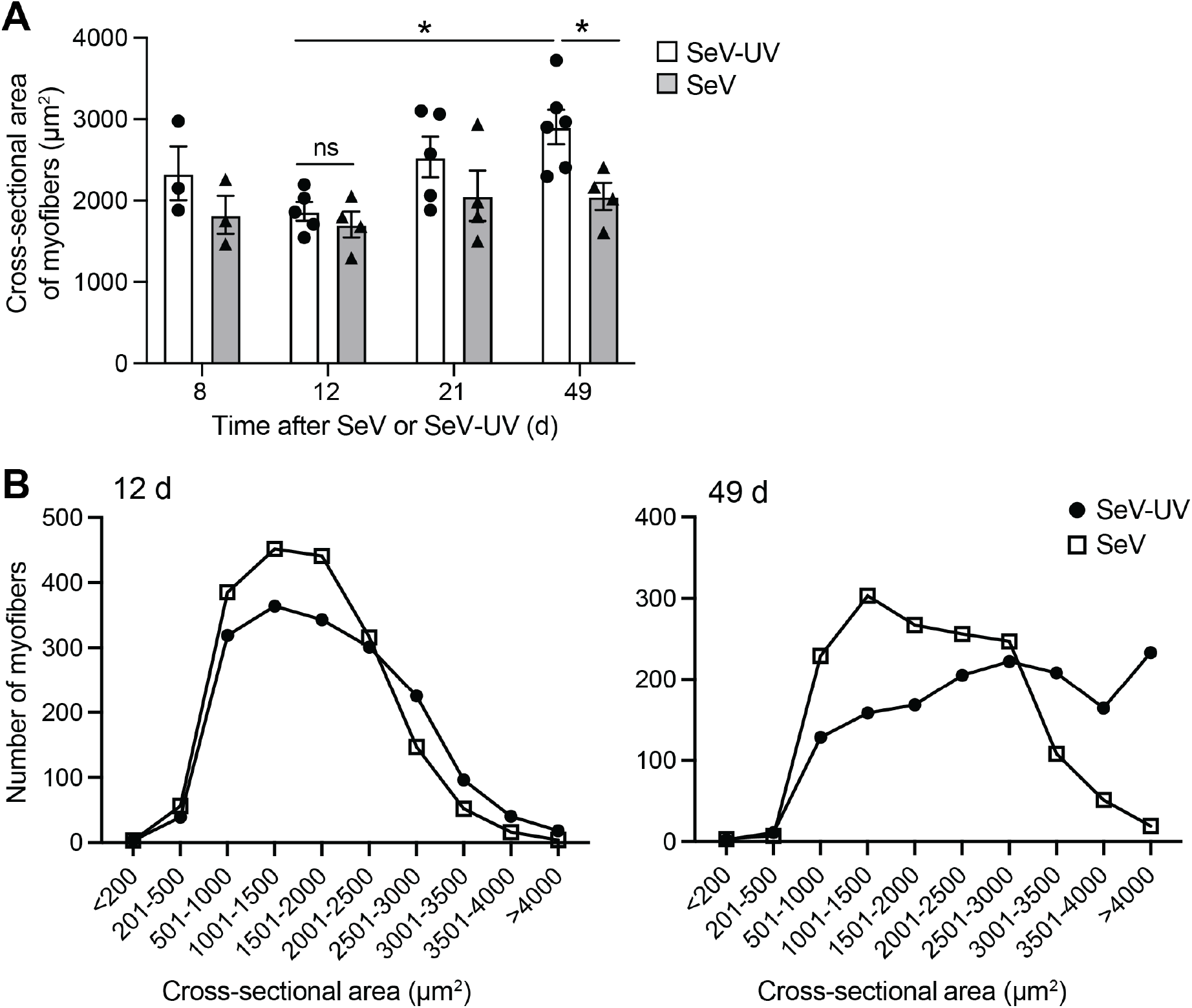
Quantitation of myofiber size in a mouse model of PVLD. A: Quantitation of myofiber cross-sectional area (μm^2^) in quadriceps muscle obtained from mice at 8, 12, 21, and 49 d after SeV infection or SeV-UV control (n = 3–6 mice per condition at each timepoint). B: Frequency distribution of myofibers from mice at 12 d and 49 d after SeV infection or SeV-UV control (n = 4-6 mice per condition at each time point). \**P*<0.05 by ANOVA.

### PVLD is associated with decreases across myofiber type

Since studies indicated a shift in myofiber type in skeletal muscle of COPD patients (7,8), the proportions of myofiber types were determined in SeV-infected and SeV-UV-control mice based on the expression of MHC isoforms in cross-sections of quadriceps muscle. For this assessment, we detected and analyzed 1,933 and 1,971 MHC^+^ myofibers in mice at 49 d after SeV infection or SeV-UV control, respectively. We found 8.5% of type I myofibers in SeV-UV mice that were undetectable in SeV-infection mice (Fig. 3A). However, all other proportions of myofiber types (i.e., type IIA, IIX and IIB myofibers) were not significantly different in SeV-infection compared to SeV-UV control conditions (Fig. 3A).

**Figure 3.**
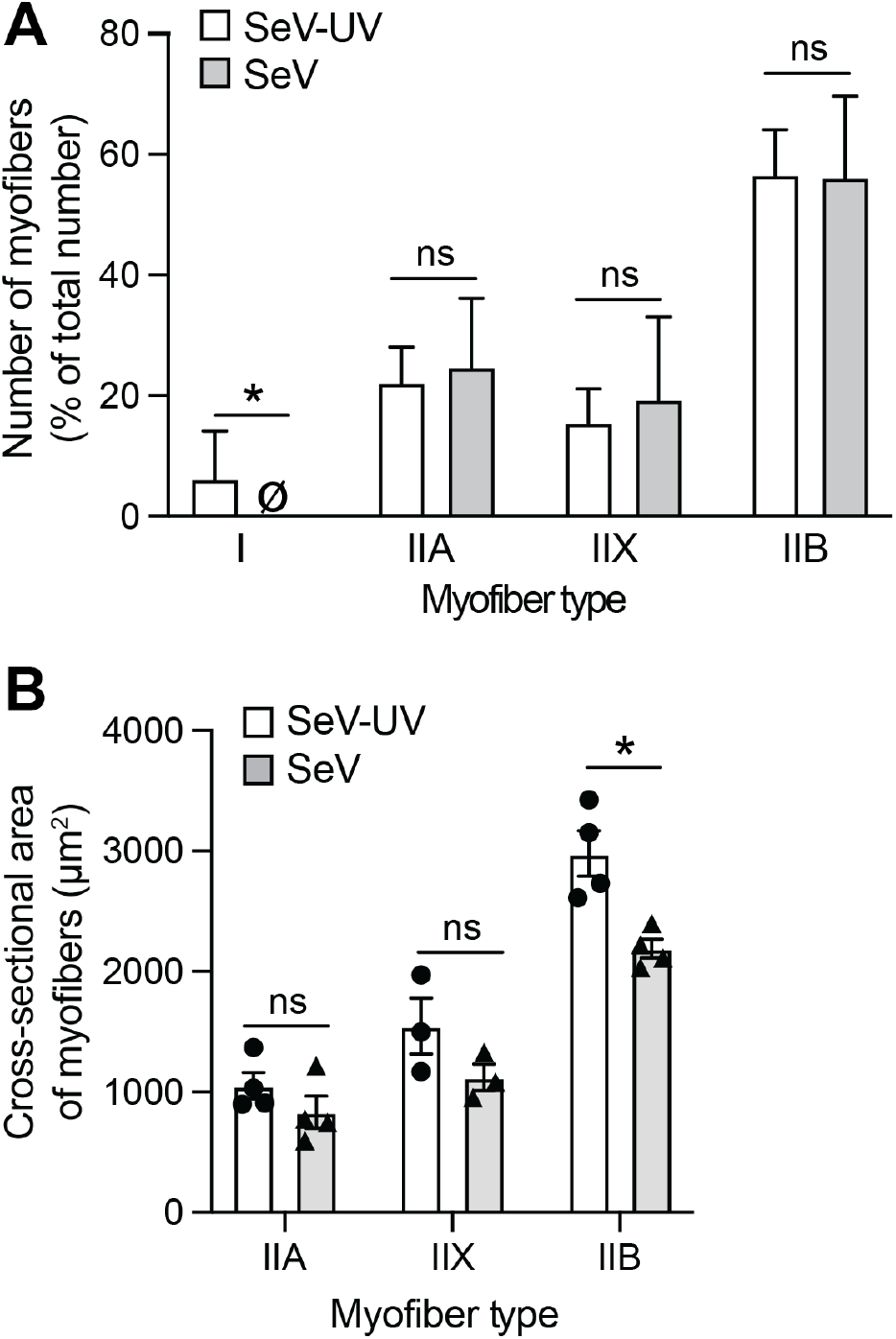
Assessment of myofiber type in a mouse model of PVLD. A: Myofiber types in quadriceps muscle obtained from mice at 49 d after SeV infection or SeV-UV control (n = 4 mice per condition). B: Myofiber cross-sectional area (μm^2^) of Type II myofibers in quadriceps muscle obtained at 49 d after SeV infection or SeV-UV control. (n = 3-4 mice per condition in each group). \**P*<0.05 by ANOVA.

We also determined cross-sectional areas of type II myofibers under the same experimental conditions. In this case, we found a selective decrease (27%) in the size of type IIB myofibers in SeV-infected compared to SeV-UV control mice with no significant change in type IIA or IIX myofibers (Fig. 3B). Correlational analysis also revealed a significant association between type IIB myofibers and total average myofiber cross-sectional area at 49 d after SeV infection (r = 0.875, p < 0.05; n = 4 mice/group). Thus, our data support a paradigm in which glycolytic myofibers are smaller despite the absence of a myofiber type shift in the SeV-infection model of PVLD.

### SeV-induced COPD is not associated with changes in ribosomal content

Our results showed a decrease in myofiber size with PVLD compared to control conditions, suggesting a possible abnormality in the ribosomal machinery needed for the synthesis of skeletal muscle proteins. Therefore, we also quantified the levels of total cellular RNA and ribosomal RNA (rRNA) during the development of PVLD in the SeV infection model. We first quantified the levels of total cellular RNA in muscle homogenates as a readout of transcriptional output (Fig. 4A). Notably, total cellular RNA relative to muscle mass was not significantly changed in relation to time or infection under these conditions (Fig. 4A). We next quantified small (18S) and large (28S) subunits of rRNA to determine ribosomal content, which is a key regulator of translation and is closely associated with regulating muscle size composing ∼80% of total cellular RNA (24). In this case, we found that no significant differences in rRNA content of skeletal muscle with respect to time or infection condition in either SeV or SeV-UV control mice (Fig. 4B).

**Figure 4.**
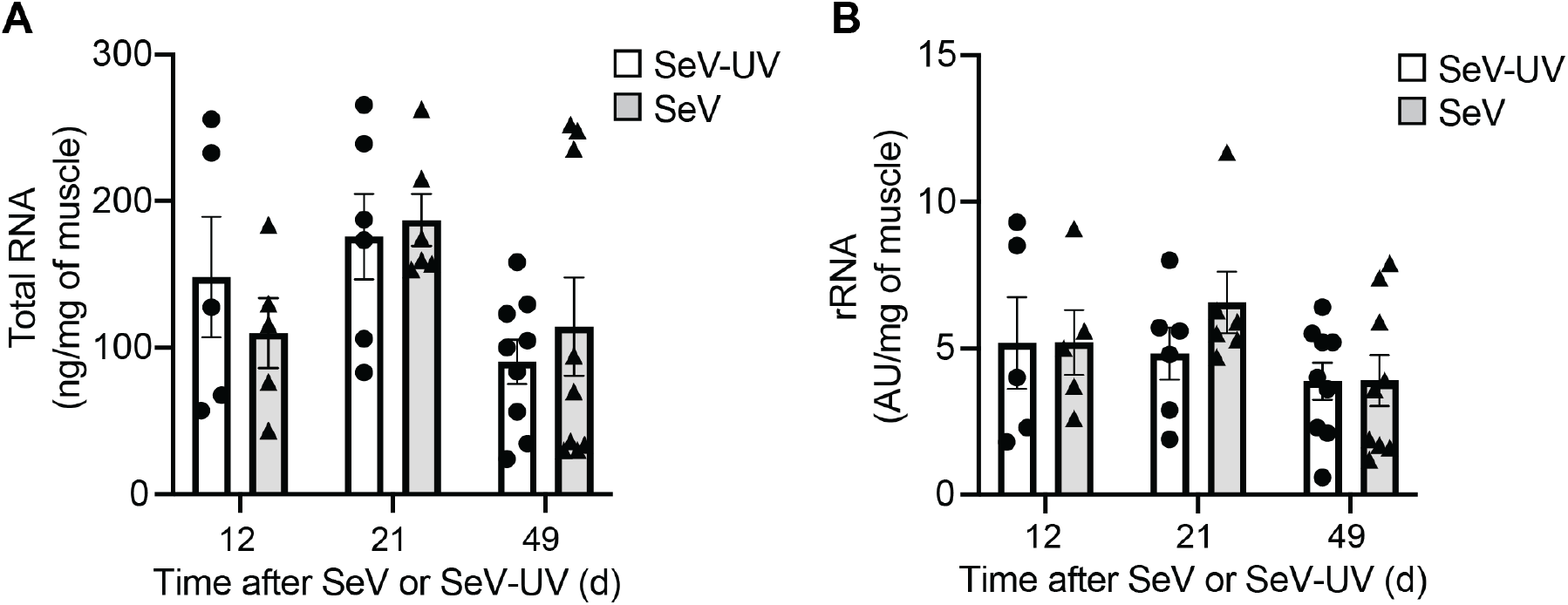
Assessment of myofiber RNA machinery a mouse model of PVLD. A: Total RNA level per mg of skeletal muscle at 12, 21, and 49 d after SeV infection or SeV-UV control (n = 5–9 mice per condition at each timepoint; \**P*<0.05 by ANOVA. B: Corresponding ribosomal RNA (rRNA) level for conditions in (A) (n = 5–9 mice per condition). ns=not significant *(P>*0.05) by ANOVA.

### PVLD is not associated with changes in the ubiquitin-proteasome system

We next tested whether the observed decreases in myofiber size might derive from increased protein degradation during our model of PVLD. We reasoned that muscle specific E3-ubiquitin ligases Fbxo32 (also known as Atrogin-1 and MAFbx) or Trim63 (also known as Murf-1) implicated in control of smooth muscle (4) could be found at increased levels after SeV infection. However, we found no significant changes in *Fbxo32* or *Trim63* mRNA at 49 d after SeV infection (Fig. 5A). Similarly, we detected no significant differences in the corresponding protein levels in skeletal muscle at 8, 12, 21, or 49 d after SeV infection compared to SeV-UV control conditions (Fig. 5B). These data suggest that the ubiquitin-proteasome pathway for degradations of skeletal muscle protein is not responsible for decreased myofiber size during the development of PVLD in the SeV mouse model.

**Figure 5.**
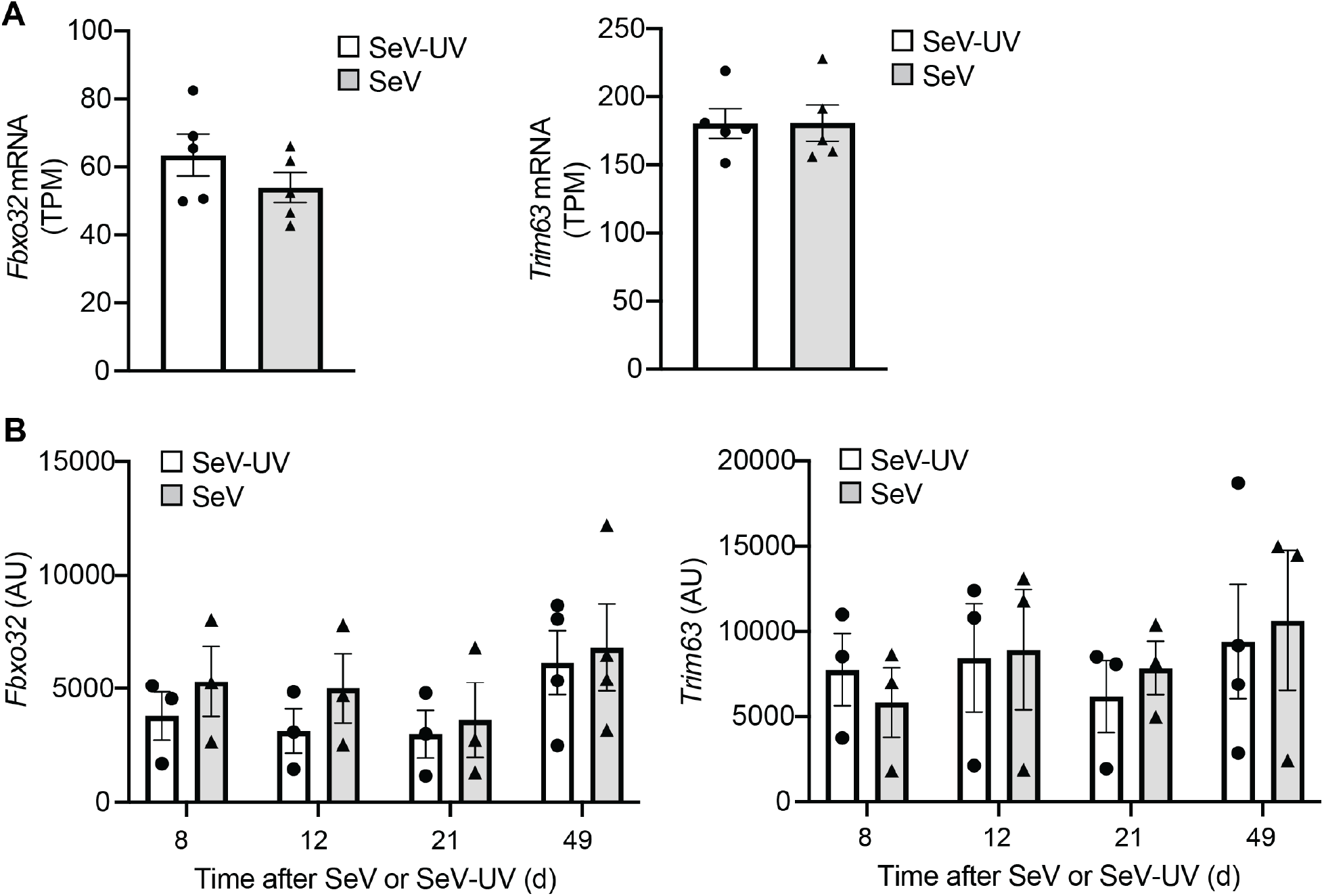
Assessment of ubiquitin proteasome levels in a mouse model of PVLD. A: Levels of *Fbxo32* and *Trim63* mRNA from RNAseq analysis at 49 d after SeV infection or SeV-UV control (n = 5 mice per condition in each group). B: Levels of Fbxo32 and Trim63 proteins at 8, 12, 21, and 49 d after SeV infection or SeV-UV control (n = 3-4 mice per condition per group). ns=not significant *(P>*0.05) by ANOVA.

## DISCUSSION

Effective treatment of chronic respiratory disease requires an improved understanding of the associated dysfunction of skeletal muscle and consequent limitation of activity. A critical barrier to studying this dysfunction is the need for experimental models of respiratory disease that also cause skeletal muscle abnormalities. The present study shows that severe respiratory viral infection in mice (using the natural pathogen SeV) causes long-term tissue remodeling not only in the lung but also in skeletal muscle. Further, the model reveals a distinct profile wherein muscle myofiber size is markedly decreased across all myofiber types but predominantly in Type IIB myofibers and is independent of gross decreases in cellular protein synthesis or degradation. Here we discuss the context and impact of these findings.

We previously established that SeV causes an acute infectious illness that transitions to long-term lung disease in a susceptible mouse strain (26, 27, 35). The model is distinct from previous approaches to study skeletal muscle dysfunction in chronic lung disease. These previous models instead primarily derive from work on COPD and therefore rely on some form of tobacco smoke exposure (30, 32). However, this approach can result in data that is difficult to interpret. Thus, smoke-induced lung disease requires continuous long-term exposure, e.g., 24 weeks or longer (15, 38), and even then might cause only mild disease (32) that is still reversible in mice (7). This result contrasts with studies of humans wherein disease persists even with cessation of tobacco smoking. Moreover, tobacco smoke exposure has a direct influence on skeletal muscle that can decrease muscle strength, capillarization, and innervation that can be detected in the absence of lung disease (11, 17). The present model thereby offers the advantages of a single trigger in time and induction of long-term lung disease and skeletal muscle dysfunction. Moreover, the model offers relevance to the currently widespread problem due to extrapulmonary manifestations of post-viral lung disease, including skeletal muscle dysfunction.

In that context, we observed a nearly 30% decrease in myofiber size in post-SeV lung disease, thereby achieving a decline that is quite similar to muscle atrophy found in COPD patients (10). Furthermore, this atrophic effect was most prominent in type IIB myofibers, wherein a 27% decrease in myofiber size indicates that it largely accounted for the overall decrease in myofiber size at the peak of PVLD. Previous work in COPD patients (21) and mouse models (17) shows that this degree of muscle atrophy results in a significant decrease in muscle strength. However, further experiments must define the loss of function in the present post-viral model. The observed decrease in myofiber size was not accompanied by any significant shift in most abundant myofiber types, in contrast to reports of a shift from oxidative to glycolytic phenotype in pulmonary disease patients (14, 19). We did detect a loss of the least abundant Type I myofibers in mice with PVLD. Whether such a myofiber shift occurs in other peripheral muscles, predominantly consisting of type I and IIA myofibers (i.e., more oxidative muscles like soleus) also warrants further investigation. Nonetheless, the present results establish a striking, persistent, and relatively selective decrease in type IIB myofiber size to support a decrease in skeletal muscle mass in association with long-term PVLD.

We also addressed the underlying pathway that might result in our observation of smaller myofibers. In that regard, we studied the abundance of rRNA as a key regulator of protein synthesis required for skeletal muscle growth (24, 25). We found that rRNA levels were not significantly different in mice with PVLD.

Further, we found no significant difference in total RNA levels suggesting a similar efficiency for RNA translation into protein despite the development of PVLD and muscle atrophy. Further studies will also need to define whether these relationships are maintained at additional time points and other muscle tissue locations. We also found no significant differences in mRNA or protein levels of muscle-specific E3 ubiquitin ligases Fbox32 and Murf-1. These factors are critical for the function of the ubiquitin-proteasome system responsible for muscle protein degradation, and increased levels and associated transcription factors were found in skeletal muscle from COPD patients (4). The present observation suggests that that PVLD limits skeletal muscle growth and myofiber size based on an alternate pathway.

Taken together, we established an experimental mouse model for investigating the impact of chronic lung disease on skeletal muscle dysfunction. This model already reveals distinct features that include a decrease in myofiber size that is relatively selective for Type IIB glycolytic myofibers and an alternative mechanism for muscle atrophy that might be independent of the usual markers of protein synthesis and degradation. Future studies of this model and related ones should address the precise pathway to the key extrapulmonary manifestation of skeletal muscle dysfunction in long-term PVLD and perhaps related forms of lung injury and remodeling disease. It will be of special interest to determine whether new therapeutic strategies from our lab and others to correct the lung remodeling disease will also impact disease phenotypes outside of the lung including skeletal muscle dysfunction.

## ACKNOWLEDGMENTS

The authors respectfully acknowledge the memory of Dr. Hughes who continues to provide a role model for scientists in this study and throughout the research community. The authors also thank the Pulmonary Morphology Core for outstanding technical support.

## Abbreviations used in this article

COPD: chronic obstructive pulmonary disease
Covid-19: coronavirus disease of 2019
MHC: myosin heavy chain
PVLD: post-viral lung disease
SARS-CoV-2: severe acute respiratory syndrome coronavirus-2
SeV: Sendai virus

## REFERENCES

1. Abdallah SJ, Voduc N, Corrales-Medina VF, McGuinty M, Pratt A, Chopra A, Law A, Garuba HA, Thavorn K, Reid RER, Lavoie KL, Crawley A, Chirinos JA, Cowan J. Symptoms, pulmonary function and functional capacity four months after COVID-19. Ann ATS in press: 2021.

2. Al-Aly Z, Xie Y and Bowe B. High-dimensional characterization of post-acute sequelae of COVID-19. Nature 594: 259–264, 2021.

3. Alevy Y, Patel AC, Romero AG, Patel DA, Tucker J, Roswit WT, Miller CA, Heier RF, Byers DE, Brett TJ, Holtzman MJ. IL-13–induced airway mucus production is attenuated by MAPK13 inhibition. J Clin Invest 122: 4555–4568, 2012.

4. Bilodeau PA, Coyne ES and Wing SS. The ubiquitin proteasome system in atrophying skeletal muscle: roles and regulation. Am J Physiol Cell Physiol 311: C392–C403, 2016.

5. Byers DE, Alexander-Brett J, Patel AC, Agapov E, Dang-Vu G, Jin X, Wu K, You Y, Alevy YG, Girard J-P, Stappenbeck TS, Patterson GA, Pierce RA, Brody SL, Holtzman MJ. Long-term IL-33-producing epithelial progenitor cells in chronic obstructive lung disease. J Clin Invest 123: 3967–3982, 2013.

6. Byers DE, Wu K, Dang-Vu G, Jin X, Agapov E, Zhang X, Battaile JT, Schechtman KB, Yusen R, Pierce RA, Holtzman MJ. Triggering receptor expressed on myeloid cells-2 (TREM-2) expression tracks with M2-like macrophage activity and disease severity in COPD. Chest 153: 77–86, 2018.

7. Caron M-A, Morissette MC, Theriault ME, Nikota JK, Stampfli MR, Debigare R. Alterations in Skeletal Muscle Cell Homeostasis in a Mouse Model of Cigarette Smoke Exposure. PloS ONE 8: e66433, 2013.

8. Centers for Disease Control and Prevention CDC. Leading causes of death, https://www.cdc.gov/nchs/fastats/leading-causes-of-death.htm. 2022.

9. del Rio C, Collins LF and Malani P. Long-term health consequences of COVID-19. JAMA 324: 1723–1724, 2020.

10. Doucet M, Russell A, Leger B, Debigare R, Joanisse DR, Caron M-A, LeBlanc P, Maltais F. Muscle atrophy and hypertrophy signaling in patients with chronic obstructive pulmonary disease. Am J Respir Crit Care Med 176: 261–269, 2007.

11. Kapchinsky S, Vuda M, Miguez K, Elkrief D, de Souza AR, Baglole CJ, Aare S, MacMillan NJ, Baril J, Rozakis P, Sonjak V, Pion C, Aubertin-Leheudre M, Morais JA, Jagoe RT, Bourbeau J, Taivassalo T, Hepple RT. Smoke-induced neuromuscular junction degeneration precedes the fibre type shift and atrophy in chronic obstructive pulmonary disease. J Physiol 596: 2865–2881, 2018.

12. Kim EY, Battaile JT, Patel AC, You Y, Agapov E, Grayson MH, Benoit LA, Byers DE, Alevy Y, Tucker J, Swanson S, Tidwell R, Tyner JW, Morton JD, Castro M, Polineni D, Patterson GA, Schwendener RA, Allard JD, Peltz G, Holtzman MJ. Persistent activation of an innate immune response translates respiratory viral infection into chronic lung disease. Nat Med 14: 633–640, 2008.

13. Kim HC, Mofarrahi M and Hussain SNA. Skeletal muscle dysfunction in patients with chronic obstructive pulmonary disease. Int J COPD 3: 637–658, 2008.

14. Lewis A, Riddoch-Contreras J, Natanek SA, Donaldson A, Man WDC, Moxham J, Hopkinson NS, Polkey MI, Kemp PR. Downregulation of the serum response factor/miR-1 axis in the quadriceps of patients with COPD. Thorax 67: 26–34, 2012.

15. Liu Q, Xu W-G, Luo Y, Han F-F, Yao X-H, Yang T-Y, Zhang Y, Pi W-F, Guo X-J. Cigarette smoke-induced skeletal muscle atrophy is associated with up-regulation of USP-19 via p38 and ERK MAPKs. J Cell Biochem 112: 2307–2316, 2011.

16. Maltais F, Decramer M, Casaburi R, Barreiro E, Burelle Y, Debigaré R, Dekhuijzen PNR, Franssen F, Gayan-Ramirez G, Gea J, Gosker HR, Gosselink R, Hayot M, Hussain SNA, Janssens W, Polkey MI, Roca J, Saey D, Schols Amwj, Spruit MA, Steiner M, Taivassalo T, Troosters T, Vogiatzis I, Wagner PD, COPD AEAHCoLMDi. An official American Thoracic Society/European Respiratory Society statement: Update on limb muscle dysfunction in chronic obstructive pulmonary disease. Am J Respir Crit Care Med 189: e15–62, 2014.

17. Nogueira L, Trisko BM, Lima-Rosa FL, Jackson J, Lund-Palau H, Yamaguchi M, Breen EC. Cigarette smoke directly impairs skeletal muscle function through capillary regression and altered myofibre calcium kinetics in mice. J Physiol 596: 2901–2916, 2018.

18. Osuchowski M, Winkler M, Skirecki T, Cajander S, Shankar-Hari M, Lachmann G, Monneret G, Venet F, Buer M, Brunkhorst F, Weis S, Garcia-Salido A, Kox M, Cavaillon J-M, Uhle F, Weigand M, Blohe S, Wiersinga W, Almansa R, de la Fuente A, Martin-LOeches I, Meisel C, Spinetti T, Schefold J, Cilloniz C, Torres A, Giamarellos-Bourboulis E, Ferrer R, Girardis M, Cossarizza A, Netea M, van der Poll T, Bermejo-Martin J, Rubia I. The COVID-19 puzzle: deciphering pathophysiology and phenotypes of a new disease entity. Lancet Respir Med 9: 622–642, 2021.

19. Puig-Vilanova E, Martínez-Llorens J, Ausin P, Roca J, Gea J, Barreiro E. Quadriceps muscle weakness and atrophy are associated with a differential epigenetic profile in advanced COPD. Clin Sci 128: 905–921, 2015.

20. Rinaldi M, Maes K, De Vleeschauwer S, Thomas D, Verbeken EK, Decramer M, Janssens W, Gayan-Ramirez GN. Long-term nose-only cigarette smoke exposure induces emphysema and mild skeletal muscle dysfunction in mice. Dis Model Mech 5: 333–341, 2012.

21. Seymour JM, Spruit MA, Hopkinson NS, Natanek SA, Man WDC, Jackson A, Gosker HR, Schols Amwj, Moxham J, Polkey MI, Wouters EFM. The prevalence of quadriceps weakness in COPD and the relationship with disease severity. Eur Respir J 36: 81–88, 2010.

22. Shiels M, Haque A, Berrington de Gonzalez A, Freedman ND. Leading causes of death in the US during the COVID-19 pandemic, March 2020 to October 2021. JaMA Internal Medicine 182: 883–886, 2022.

23. Swallow EB, Reyes D, Hopkinson NS, Man WDC, Porcher R, Cetti EJ, Moore AJ, Moxham J, Polxkey MI. Quadriceps strength predicts mortality in patients with moderate to severe chronic obstructive pulmonary disease. Thorax 62: 115–120, 2007.

24. von Walden F, Casagrande V, Östlund Farrants A-K, Nader GA. Mechanical loading induces the expression of a Pol I regulon at the onset of skeletal muscle hypertrophy. Am J Physiol Cell Physiol 302: C1523–C1530, 2012.

25. von Walden F, Liu C, Aurigemma N, Nader GA. mTOR signaling regulates myotube hypertrophy by modulating protein synthesis, rDNA transcription, and chromatin remodeling. Am J Physiol Cell Physiol 311: C663–C672, 2016.

26. Walter MJ, Kajiwara N, Karanja P, Castro M, Holtzman MJ. IL-12 p40 production by barrier epithelial cells during airway inflammation. J Exp Med 193: 339–352, 2001.

27. Walter MJ, Morton JD, Kajiwara N, Agapov E, Holtzman MJ. Viral induction of a chronic asthma phenotype and genetic segregation from the acute response. J Clin Invest 110: 165–175, 2002.

28. Wang X, Wu K, Keeler SP, Mao D, Agapov EV, Zhang Y, Holtzman MJ. TLR3-activated monocyte-derived dendritic cells trigger progression from acute viral infection to chronic disease in the lung. J Immunol 206: 1297–1314 (selected for Top-Reads p. 1115), 2021.

29. Watz H, Waschki B, Boehme C, Claussen M, Meyer T, Magnussen H. Extrapulmonary Effects of Chronic Obstructive Pulmonary Disease on Physical Activity. Am J Respir Crit Care Med 177: 743–751, 2008.

30. Wheaton AG, Liu Y, Croft JB, VanFrank B, Croxton TL, Punturieri A, Postow L, Greenlund KJ. Chronic obstructive pulmonary disease and smoking status—United States, 2017. MMWR Morb Mortal Wkly Rep 68: 533–538, 2019.

31. World Health Organization WHO. The top 10 causes of death, https://www.who.int/news-room/fact-sheets/detail/the-top-10-causes-of-death. 2020.

32. Wright JL, Cosio M and Churg A. Animal models of chronic obstructive pulmonary disease. Am J Physiol Lung Cell Mol Physiol 295: L1–L15, 2008.

33. Wright JL and Sun JP. Effect of smoking cessation on pulmonary and cardiovascular function and structure: analysis of guinea pig model. J Appl Physiol 76: 2163–2168, 1994.

34. Wu K, Byers DE, Jin X, Agapov E, Alexander-Brett J, Patel AC, Cella M, Gilfilan S, Colonna M, Kober DL, Brett TJ, Holtzman MJ. TREM-2 promotes macrophage survival and lung disease after respiratory viral infection. J Exp Med 212: 681–697, 2015.

35. Wu K, Kamimoto K, Zhang Y, Yang K, Keeler SP, Gerovac BJ, Agapov EV, Austin SP, Yantis J, Gissy KA, Byers DE, Alexander-Brett J, Hoffmann CM, Wallace M, Hughes ME, Morris SA, Holtzman MJ. Basal-epithelial stem cells cross an alarmin checkpoint for post-viral lung disease. J Clin Invest 131: e149336, 2021.

36. Wu K, Wang X, Keeler SP, Gerovac BJ, Agapov E, Byers DE, Gilfillan S, Colonna M, Zhang Y, Holtzman MJ. Group 2 innate lymphoid cells must partner with the myeloid-macrophage lineage for long-term postviral lung disease. J Immunol 205: 1084–1101, 2020.

37. Wu K, Zhang Y, Yin Declue H, Austin SR, Byers DE, Crouch EC, Holtzman MJ. Lung remodeling regions in long-term Covid-19 feature basal epithelial cell reprogramming. medRxiv 10.1101/2022.09.17.22280043: 2022.

38. Xiong J, Le Y, Rao Y, Zhou L, Hu Y, Guo S, Sun Y. RANKL Mediates Muscle Atrophy and Dysfunction in a Cigarette Smoke-induced Model of COPD. Am J Respir Cell Mol Biol 64: 617–628, 2021.

39. Zhang Y, Mao D, Keeler SP, Wang X, Wu K, Gerovac BJ, Shornick LP, Agapov E, Holtzman MJ. Respiratory enterovirus (like parainfluenza virus) can cause chronic lung disease if protection by airway epithelial STAT1 is lost. J Immunol 202: 2332–2347, 2019.

